# Functional Brain Mapping of Body Size Estimation Using a 3D Avatar

**DOI:** 10.1101/2025.07.24.666647

**Authors:** Hayden J. Peel, Joel P. Diaz-Fong, Sameena Karsan, Rajay Kumar, Gerhard Hellemann, Jamie D. Feusner

## Abstract

**Background:** Body size estimation—the ability to judge the size and shape of one’s own body— is a key perceptual component of body image. However, its neural basis, and the basis for inter-individual differences in accuracy, remain poorly understood, partly due to limitations in existing assessment tools.

**Methods:** We developed Somatomap 3D, an interactive fMRI-compatible task allowing participants to manipulate a rotatable 3D avatar by adjusting the size and shape of 26 individual body parts to match their perceived body. Twenty-eight healthy male and female adults completed the task during fMRI. Brain activity in a priori regions of interest from previous studies of body processing was modeled using a general linear model incorporating event-specific parameters and parametric modulators related to task performance. Inter-individual differences in body size estimation accuracy were calculated using multidimensional scaling of body part estimation errors, and scores were correlated with BOLD signal eigenvariates from regions of interest.

**Results:** Task engagement was associated with significant activation in hypothesized body-selective and multisensory regions, including bilateral extrastriate body area, right fusiform body area, right superior parietal lobule, and bilateral premotor cortex. Multidimensional scaling identified a primary subdimension reflecting distortions in body part girths, which was significantly associated with neural responses in the superior parietal lobule. No other brain regions showed significant associations with inter-individual differences in estimation accuracy.

**Conclusions:** These results suggest that body size estimation engages a distributed network of visual, motor, and parietal regions. Among these, only the superior parietal lobule showed a significant association with inter-individual variation in body size estimation accuracy for body part girths, supporting its role as a candidate neural substrate for altered body representation in psychiatric conditions such as eating disorders and body dysmorphic disorder.

## Introduction

Perception of one’s body is a multidimensional construct shaped by visual, somatosensory, proprioceptive, and vestibular systems, integrated across distributed brain networks (Feusner & Cazzato, n.d.). A key perceptual component of body image is body size estimation (BSE)—the ability to visually judge the size and shape of one’s own body (Cash & Deagle III, 1997; Farrell et al., 2005; Gardner & Brown, 2014). Disruption in BSE is common in psychiatric conditions such as eating disorders (e.g., anorexia nervosa) (Ralph-Nearman et al., 2021) and body dysmorphic disorder (Karsan et al., 2024). Yet, despite its clinical relevance, current tools to assess BSE often lack ecological validity and spatial specificity, and the neural mechanisms supporting BSE remain incompletely understood.

BSE tasks typically use metric and depictive techniques (Mölbert et al., 2017). Metric methods involve direct size estimations using tools such as tape measures or calipers. Depictive methods—more commonly used—require participants to compare their perceived body size to visual representations, such as body drawings (Uher et al., 2005) or size-modified silhouettes (Hamamoto et al., 2023). While informative, these methods typically use static, two-dimensional stimuli that lack the depth, orientation, and perspective of naturalistic visual body perception.

Other paradigms using digitally distorted photographs [e.g., (Castellini et al., 2013; Miyake et al., 2010) or virtual avatar comparisons [e.g., (Gao et al., 2016; Owens et al., 2010) offer improved visual realism and body shape control, but often remain static and lack participant interaction. Interactive 3D approaches, such as manipulable avatars, allow participants to actively engage with a dynamic body model, better approximating real-world self-perception like viewing oneself rotating in front of a mirror. If the goal is to understand how visual processing of the body may be altered in conditions involving body image disturbance, it is essential to use stimuli that engage the visual system in a more dynamic manner.

To address these limitations, we developed Somatomap 3D, a novel tool that allows participants to interactively rotate and manipulate a 3D avatar by adjusting the size and shape of 26 body parts to match their perceived current body (Ralph-Nearman et al., 2019). Unlike prior avatar-based paradigms that algorithmically applied fat distributions that scale larger or smaller across the whole body according to population averages (Cornelissen et al., 2017; Mölbert et al., 2017), Somatomap 3D enables localized estimation and adjustment of body parts. This feature is important given BSE distortions in conditions like anorexia nervosa often affect specific areas such as the abdomen, hips, or thighs (Ralph-Nearman et al., 2021). Prior studies using Somatomap have identified body part specific over-and under-estimations in nonclinical samples (Ralph-Nearman et al., 2019) and in clinical samples of individuals with body dysmorphic disorder and anorexia nervosa (Karsan et al., 2024; Ralph-Nearman et al., 2021), and shows high test-retest reliability (Ralph-Nearman et al., 2023). However, its neural correlates remain unknown. To date, no study has combined body-part-specific 3D avatar manipulation with neuroimaging to investigate the neural basis of BSE. Identifying these mechanisms is essential for refining models of body perception and informing interventions for body image disturbances.

Prior research has implicated several brain regions in visual body processing and possibly BSE. These include the extrastriate and fusiform body areas (EBA and FBA), which show selective responses to human bodies and are implicated in body part recognition, self-referential processing, and the perception of body size and shape (Downing & Peelen, 2011; Hummel et al., 2013; Urgesi et al., 2007; Vocks et al., 2010). Also involved in body processing are the temporo-parietal junction (TPJ), a hub for multisensory integration, spatial perspective-taking, and body ownership (Blanke et al., 2005; Castellini et al., 2013); the superior parietal lobule (SPL), associated with visuospatial attention and metric body representation (Castellini et al., 2013; Gaudio & Quattrocchi, 2012); the premotor cortex (PMC), given its role in motor imagery and mental body transformations (Gao et al., 2016; Haggard, 2008; Hanakawa et al., 2008); and the primary visual cortex (V1), given its role in early visual processing and size perception (Schwarzkopf et al., 2011). Together, these regions formed the basis of our *a priori* regions of interest (ROI), selected based on theoretical and empirical links to perceptual and visuospatial aspects of the Somatomap 3D task.

The goal of this study was to characterize neural activation patterns associated with engagement in BSE, and those associated with its accuracy, using fMRI. We hypothesized that the EBA, FBA, TPJ, SPL, PMC, and V1 would be engaged during avatar manipulation, and that neural activity would scale with trial-by-trial variation in body part estimation accuracy.

Accordingly, we performed parametric modulation analyses within the ROIs using *BSE percent error*, the trial-wise percent discrepancy between estimated and actual body part size, divided by the actual body part size, as a continuous regressor. We also applied multidimensional scaling (MDS) to participants’ estimation error profiles across the 26 body parts. This unsupervised machine learning approach was motivated by the goal of characterizing inter-individual differences in how people perceive and represent their own bodies. MDS allowed us to reduce data complexity by summarizing participants’ perceptual profiles into a smaller number of latent dimensions that reflect systematic patterns in estimation errors across the body. These data-driven dimensions provide a compact, multivariate representation of BSE performance that captures how each individual’s pattern of errors is positioned within the structure of estimation patterns identified across participants. We then tested how inter-individual differences along these dimensions are associated with BSE-related neural activation in the ROIs. We hypothesized that differences in participants’ overall patterns of body part estimation errors would be associated with activation in the same ROIs during body estimation. We also conducted whole-brain event-related analyses and parametric modulation to explore broader neural responses related to task features. Together, these approaches aimed to map the functional neural activation patterns underlying BSE and those associated with BSE accuracy.

## Methods

### Participants

This study was approved by the Centre for Addiction and Mental Health (CAMH) Research Ethics Board (075-2021, 2022). Thirty participants (12 males, mean age = 24.43 (SD ± 5.15), age range = 18-40, mean years of education = 15.83, SD ± 3.36)) from the Greater Toronto Area were recruited as part of a larger study of visual processing. Participants met with a clinician who performed a structured diagnostic interview and were excluded if they met criteria for any current DSM-5 disorders, neurological disorder, or any major medical disorder that may affect cerebral metabolism (e.g., diabetes, thyroid disorders), were pregnant, could not safely be scanned (e.g., ferromagnetic implants or other devices), or had visual acuity (corrected or uncorrected) worse than 20/35 in each eye as verified with a Snellen close vision eye chart. All provided informed consent prior to enrolment. Two participants were excluded due to technical errors during data acquisition. The final analyzed sample comprised 28 participants (12 males and 16 females, mean age = 24.25 (SD ± 5.28), age range = 18-40, mean years of education = 15.61 (SD ± 3.36)).

## Procedures

### Somatomap 3D task

Participants used *Somatomap 3D* to engage with a three-dimensional avatar on a computer screen, as previously described (Karsan et al., 2024; Ralph-Nearman et al., 2019, 2021).

Participants used sliders to adjust 26 individual body parts on a rotatable 3D avatar to reflect their perceived current body shape (see Figure 1). They were instructed to “configure the body (from the neck down) to best match your current body shape.” Adjustments were made one body part at a time, and participants were allowed to revisit and modify any previously adjusted parts until satisfied with the overall configuration. The body parts included: neck length and girth, shoulder width, bust girth, chest girth, abdomen protrusion, upper arm length and girth, elbow girth, lower arm length and girth, wrist girth, hand size and length, torso length, waist size, hip size and width, thigh length and girth, knee girth, lower leg length and girth, ankle girth, and feet width and length. Participants were given 10 minutes to complete the task and were given a warning when they had 3 minutes remaining.

**Figure 1.**
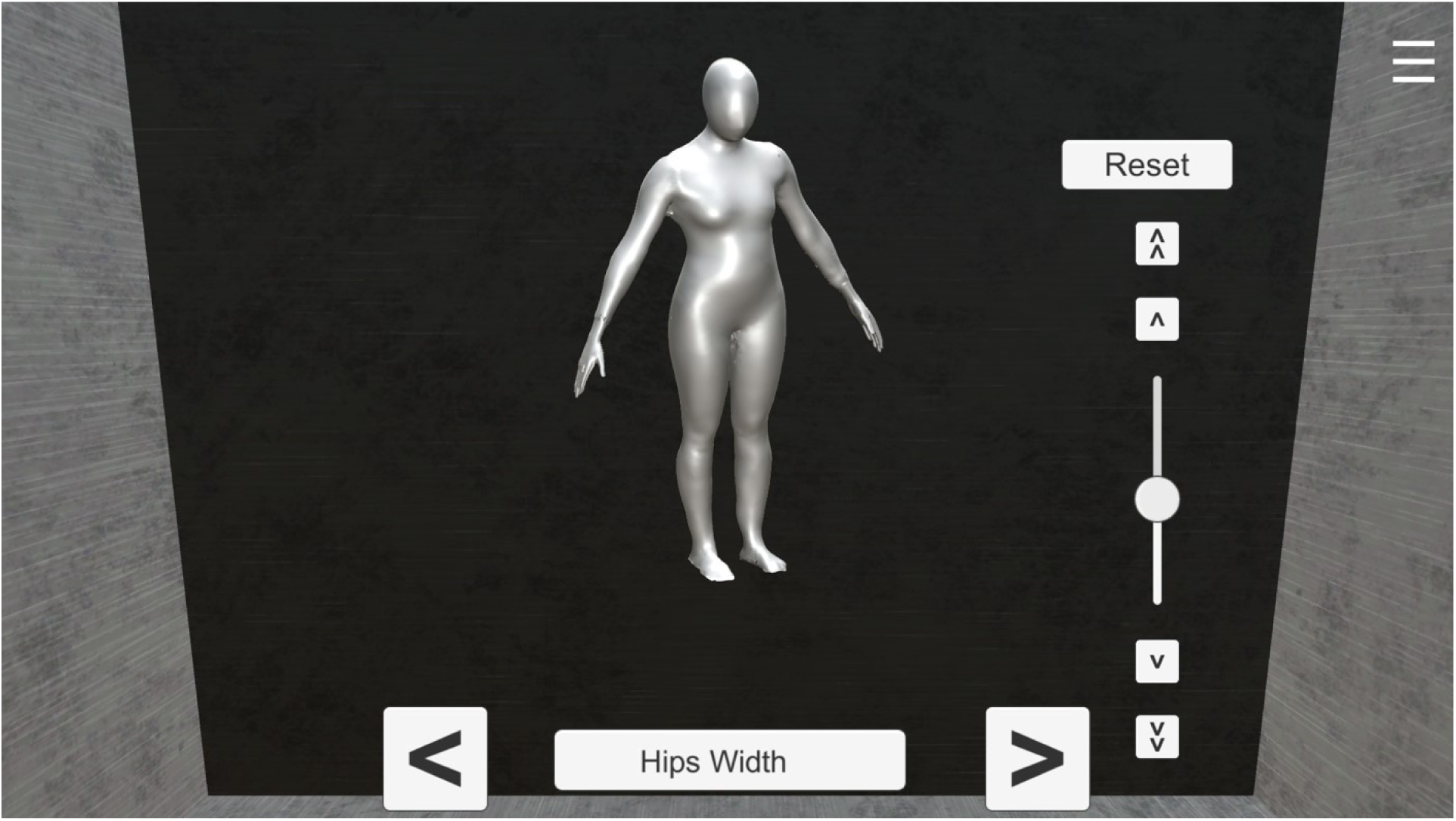
Example of the Somatomap 3D interface. Participants selected either a masculine-appearing or feminine-appearing avatar and could adjust 26 different body parts using the slider located on the right. Body parts were navigated using the left and right buttons on the bottom panel, which corresponded to the previous and next body part, respectively. The currently selected body part was displayed centrally. Avatars could be freely rotated along the x-and y-axes to allow inspection and modification from multiple viewing angles.

Participants first completed the task on a desktop computer to get accustomed to using a trackball, and then performed the task again in the scanner. All data presented here corresponds to the scanner session, completed using an MR-compatible trackball (Current Designs, Inc., Philadelphia, United States). For this study, the *Somatomap 3D* task was packaged in Unity (v2021.1.5f1; Unity Technologies, San Franscisco, CA, United States) as a standalone Windows Desktop application. The WebLink software (v2.2.161; SR Research Ltd., Ottowa, ON, Canada) was utilized to capture screen recordings, eye movements, and mouse and keyboard events, including the fMRI triggers used to synchronize events with the scan (see Supplementary Materials S1 for more details).

### Body measurements

Research staff obtained physical measurements of the corresponding 26 body parts after the *Somatomap 3D* fMRI task procedures. Physical measurements were taken using a tape measure and were completed twice for the left and right side (where applicable) and then averaged. See Groe et al. (2025) for a detailed description of how each body part was measured. Height and weight measurements were recorded using a stadiometer and a calibrated scale, respectively.

## Data Acquisition

The MRI data were acquired using a Siemens MAGNETOM Prisma 3T MRI scanner (Siemens Healthineers, Erlangen, Germany) at the Toronto Neuroimaging Facility located at the University of Toronto. Imaging was conducted with a 32-channel head coil. Structural images were acquired using a T1-weighted MPRAGE sequence (TR/TE: 2300/2.27 ms; flip angle: 8°; 256 x 256 matrix; voxel size: 1 mm^3^; 192 slices). Functional images were acquired using a multi-band echo planar imaging (EPI) sequence (TR/TE: 1000/30 ms; multi-band acceleration factor: 5; flip angle: 60°; 104 x 104 matrix; voxel size: 2 mm³; 62 slices). Additionally, field maps were acquired in opposite phase encoding directions as echo planar spin-echo (epse) sequences (TR/TE: 6629/60 ms; flip angle: 90°; 104 x 104 matrix; voxel size: 2 mm^3^; 65 slices) to estimate the displacement map for susceptibility distortion correction.

### Preprocessing

Images were processed using *fMRIPrep* v22.0.2 (Esteban et al., 2019). For more detailed information about the fMRI preprocessing, please refer to the Supplementary Materials S2. Briefly, spatial normalization of the T1-weighted image to standard MNI space was performed through nonlinear registration. The processing of the BOLD timeseries consisted of head-motion estimation, slice time correction, and susceptibility distortion correction utilizing two spin echo field maps of opposite phase encoding directions. The processed BOLD timeseries were then resampled in their native space in a single interpolation step, and finally, resampled into standard MNI space, generating the spatially normalized, preprocessed BOLD runs. Spatial smoothing was performed with a Gaussian kernel of 6 mm FWHM prior to automated removal of motion artifacts with independent component analysis (ICA-AROMA; Pruim et al., 2015). Volumes with excessive motion, as determined by DVARS from the FSL motion outliers’ tool (with default settings), were included as confound regressors in the first-level analyses described below.

### Mask Creation

ROIs were defined using term-based meta-analytic activation maps from Neurosynth (www.neurosynth.org), further constrained using probabilistic anatomical masks from the Harvard-Oxford and Jülich atlases to improve anatomical specificity. Body-selective visual areas were defined as the overlap between the “body” Neurosynth map and the Harvard-Oxford 50% probabilistic maps for the a) Inferior Lateral Occipital Cortex for the EBA, and the b) Posterior Inferior Temporal Gyrus and Posterior Temporal Fusiform Cortex for the FBA. Note, this yielded bilateral EBA, but only right-lateralized FBA. Bilateral TPJ was defined as the overlap between the Neurosynth map for the term “tpj” and the Harvard-Oxford Angular Gyrus mask. Bilateral PMC was defined as the intersection between the Neurosynth map for “premotor cortex” and the Jülich probabilistic maps for BA6 in the left and right hemispheres. Bilateral SPL was defined by the overlap of the Neurosynth map for “superior parietal” and six Jülich probabilistic maps: areas 7A, 7P, and 7PC in both hemispheres. Bilateral V1 was generated by intersecting the Neurosynth map for “primary visual” with the Jülich probabilistic V1 maps from both hemispheres. All six ROIs were then merged into a single mask for subsequent analyses (see *Figure 2*).

**Figure 2.**
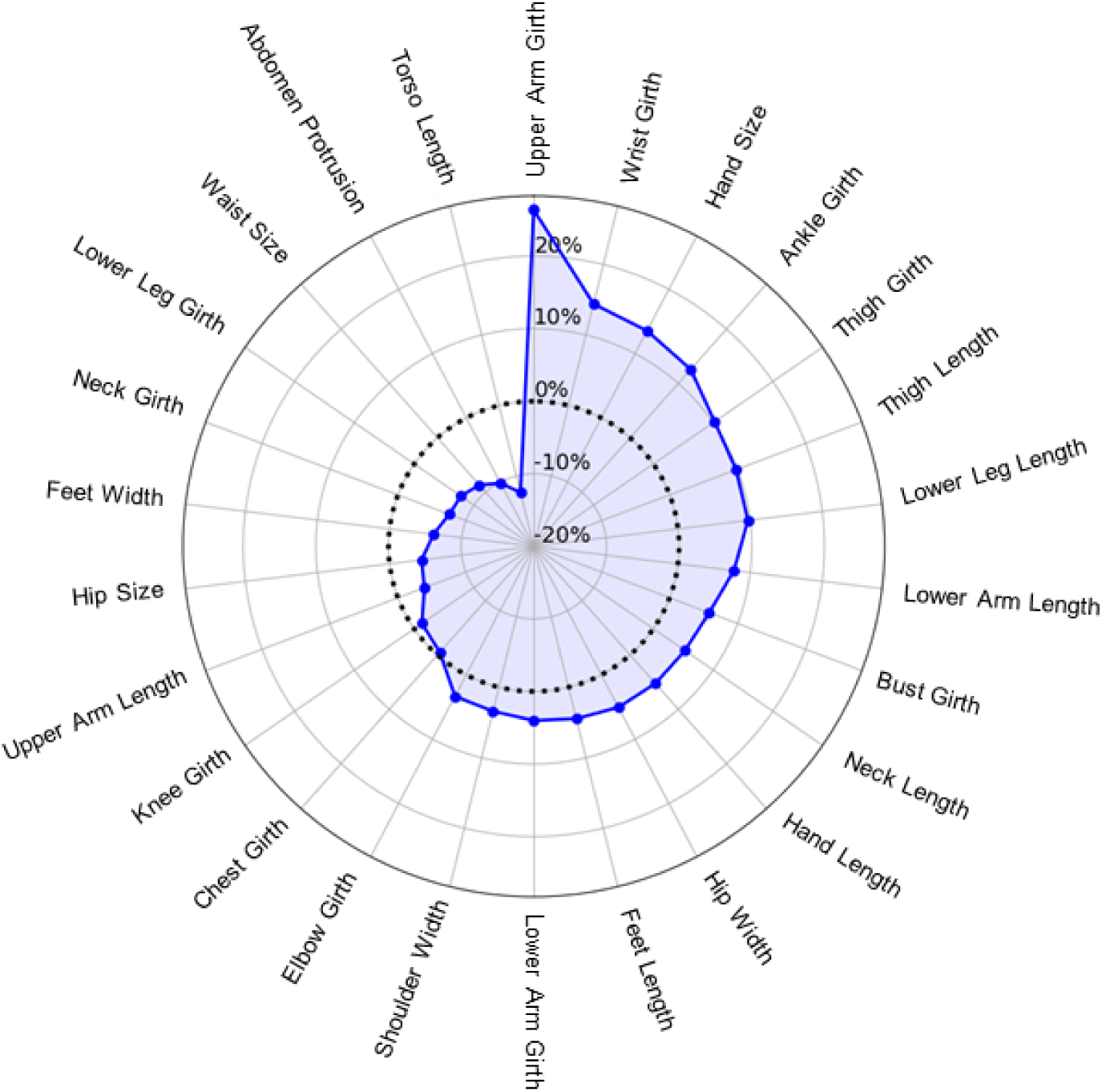
Radar chart depicting mean body size estimation (BSE) inaccuracy (percent error) across 26 body parts, arranged clockwise in the order of overestimation to underestimation. Positive values indicate mean overestimation of body part size, negative values indicate mean underestimation, and zero (dotted line) denotes perfect accuracy.

## Data Analysis

### BSE measurements

Body measurements from *Somatomap 3D* were converted into centimetres from arbitrary units using previously described methods (Ralph-Nearman et al., 2019). For each body part, a BSE percent error score was calculated by subtracting the actual physical measurement from one’s estimated body part size obtained from *Somatomap* and dividing by the actual measurement.

This method allows for direct comparisons across individuals and body parts of varying sizes by expressing estimation error relative to the actual measurement. This is desirable given some body parts are larger, and thus, are prone to larger deviations in raw centimetre error according to general psychophysical principles like Weber’s Law (Fechner, 1966).

### fMRI Modeling and Within-Mask Analyses

Using mouse click timestamps, *BSE size adjustment events* were defined in the time-series as the periods when the participant was adjusting the avatar body size. *Rotation events* were defined in the time-series as periods when the participant was rotating the position of the avatar. The contrast included intervening periods between these rotations and size adjustments, when participants used the trackball to navigate between body parts (see Supplementary Materials S1 for additional details). These intervals control for low-level motor activity and visuospatial processing demands associated with body part selection, isolating perceptual and decisional components of the BSE task.

Preprocessed fMRI data were analyzed using FSL FEAT (v6.0.4). At the first level, a single general linear model (GLM) was constructed that included all task events and several parametric regressors. Events included (i) *BSE size adjustment* and (ii) *rotation events,* both modeled with durations equal to the event-specific response time (RT), which were combined at the second level to construct the (iii) *‘avatar engagement’ events*. Our primary interpretive focus was on the *avatar engagement* periods, as the self-paced, continuous nature of Somatomap 3D often results in BSE and rotation processes unfolding in close succession, making a combination of the two an integrated measure of task engagement. Parametric regressors included (i) *BSE size adjustment events* modulated by the demeaned BSE percent error for the event (ii) *BSE size adjustment events* modulated by the demeaned RT, and (iii) *rotation events* modulated by the demeaned RT. All parametric regressors were modeled as event-related and with durations matched to trial-specific RT. The BSE percent error regressor was included in line with our hypotheses and research questions, while RT-based parametric regressors were included to account for variability in trial duration. At the group level, analyses were conducted using FSL’s FLAME 1 (FMRIB’s Local Analysis of Mixed Effects, stage 1), with sex included as a covariate to account for differences in the avatar model across sexes. (All participants were presented with the avatar model that aligned with sex assigned at birth.) Within-mask analyses constituted our primary analytic approach. Contrast estimates from the first-level GLM were entered into the group-level analysis, and statistical inference was performed using a cluster-based thresholding approach, with clusters defined at Z > 3.1 and a family-wise error (FWE)-corrected cluster significance threshold of p < 0.05 applied across the entire mask to control for multiple comparisons.

### Multidimensional scaling (MDS) of BSE accuracy calculations

To capture inter-individual differences in BSE across 26 body parts in a parsimonious manner, we applied MDS, an unsupervised, data-driven machine learning technique that identifies latent dimensions underlying latent patterns of dissimilarity—in this case, differences in BSE percent error—by positioning each participant in a low-dimensional space based on their distance from others. We used the smacof package in R (Leeuw & Mair, 2009; Mair et al., 2022). The smacof algorithm was chosen for its flexible implementation and reliable convergence properties, offering improved stability and fit compared to classical MDS algorithms (de Leeuw, 1977).

Dissimilarity was quantified as pairwise Euclidean distances across body part errors, and the MDS algorithm positioned participants to best preserve these distances in the output space. This yielded latent dimensions that captured dominant patterns of variation in BSE distortion. Prior to calculating the MDS dimensions, participants’ height, weight, body mass index, and gender were regressed out of percent error scores to control confounding effects. MDS was then conducted on the residuals, producing participant coordinates along each dimension.

### Associations between MDS and BSE-related brain activation

To test whether inter-individual differences in BSE accuracy were associated with BSE-sensitive neural responses, we derived subject-level, ROI-specific estimates of task-related activation to serve as input for correlational analyses with MDS subdimension scores. We extracted the mean of the first principal eigenvariate of the BOLD time series associated with BSE events from voxels within each ROI, separately. This measure captures the dominant pattern of task-related signal change within each ROI, providing a summary index of activation intensity across voxels during body estimation. We selected the eigenvariate approach because alternatives such as percentage signal change are based on homogenous averaging across voxels (Friston et al., 2006), which could “dilute” the signal of interest when an ROI contains both activated and deactivated voxels. Pearson correlations were performed between MDS subdimension scores and eigenvalues, with the false discovery rate (FDR) adjusted using the Benjamini-Hochberg procedure for six comparisons (FBA, EBA, SPL, PMC, TPJ, and V1).

### Exploratory Whole-Brain Analyses

Complementing the within-mask analyses, we conducted an exploratory whole-brain analysis using the same regressors described above. Group-level statistical maps were generated with the same cluster-based correction (Z > 3.1; FWE p <.05), applied across the entire brain. These results were interpreted descriptively to identify additional task-related regions outside the *a priori* ROIs.

## Results

### Behavioural Results

Across all participants’ BSE trials (n = 2,852 total), the mean trial duration was 3.28 seconds (SD = 2.43), with a median of 2.68 seconds and an interquartile range (IQR) of 2.64 seconds. Interstimulus interval (ISI) durations were defined as the time elapsed between consecutive events. The mean ISI duration was 4.14 seconds (SD = 3.24), with a median of 3.82 seconds and an interquartile range (IQR) of 3.47 seconds. For further details concerning behavioural performance, see Supplementary Materials S3. As can be visualized in radar plots (see *Figure 2*), participants tended to overestimate some body parts (16/26), and underestimated others (10/26).

### Within-Mask Results

The ROI analysis showed significant activation for avatar engagement in the right FBA, right SPL, bilateral PMC, and bilateral EBA (see *Table 1* and *Figure 3*). No significant activity was detected in the TPJ or V1.

**Figure 3.**
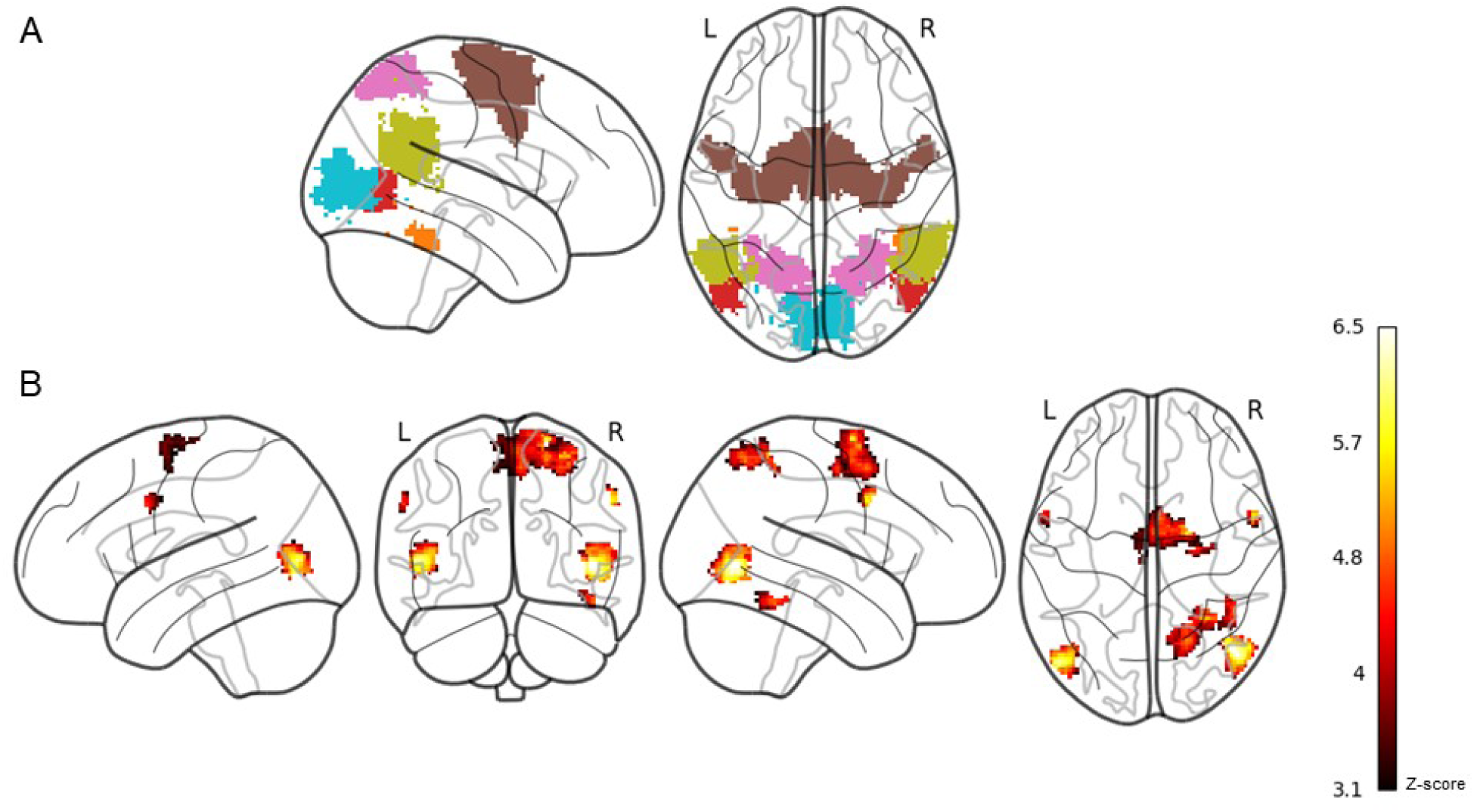
Statistical maps showing significant activation clusters and ROI mask visualizations. Panel (A) Colour-coded ROI mask: red = extrastriate body area, orange = fusiform body area, blue = primary visual cortex, gold = temporoparietal junction, pink = superior parietal lobule, brown = premotor cortex. These regions were merged into a single mask for the ROI analysis. Panel (B) Regions showing increased BOLD response during avatar engagement, thresholded at *z* > 3.1. A dark red-to-yellow colour scale indicates increasing *z*-score magnitude. Maps are displayed on a glass brain.

**Table 1.** Clusters peaks associated with ROI-specific avatar engagement analyses.

| Cluster Peak | Size (voxels) | Z max | MNI Coordinates |  |  |
| --- | --- | --- | --- | --- | --- |
|  |  |  | *X* | *Y* | *Z* |
| R PMC | 719 | 5.75 | 20 | -4 | 70 |
| R EBA | 457 | 6.63 | 46 | -68 | 2 |
| R SPL | 382 | 5.27 | 30 | -50 | 62 |
| L EBA | 272 | 6.16 | -50 | -74 | 4 |
| R FBA | 75 | 4.84 | 46 | -46 | 20 |
| R PMC | 42 | 4.57 | 26 | -12 | 54 |
| L PMC | 36 | 4.54 | -58 | 8 | 34 |
| R PMC | 35 | 5.63 | 56 | 4 | 36 |

*L* left hemisphere, *R* right hemisphere, *EBA* Extrastriate Body Area, *PMC* premotor cortex, *SPL* superior parietal lobule, *FBA* fusiform body area, *MNI* Montreal Neurological Institute. $Z > 3.1$ , cluster-corrected $p < .05$ FWE

Parametric modulation analyses within the ROI mask showed no significant effects for BSE percent error. Results from other event-related and parametric regressors are reported in the Supplement (see Supplementary Materials S4).

## MDS Results

A three-dimensional MDS solution was selected to balance model fit and interpretability. Stress values decreased from 0.40 in one dimension to 0.25 in two dimensions and 0.17 in three, with further reductions tapering off thereafter (e.g., stress = 0.13 in 4D, 0.10 in 5D), supporting a three-dimension solution as a reasonable approximation (see Supplementary Figure S6). We interpreted the three dimensions by computing Pearson correlations between each participant’s subdimension scores and their estimation error for each body part. Subdimension 1 was correlated with errors for girth-related body parts across the avatar (e.g., neck girth, hip size, arm girth etc.; *r* range.30 to.86) (see Supplementary Figure S7). In contrast, subdimensions 2 and 3 showed more heterogeneous patterns, with moderate correlations involving both length and height-based body parts. Given its relevance for potential future clinical applications in eating disorders and body dysmorphic disorder, often characterized by concerns about body parts girths being too large (i.e. “fat”), we used subdimension 1 for the analysis with BSE-related brain activation.

### MDS Associations with BSE-related Brain Activation Results

A significant negative correlation was detected between right SPL activity and MDS subdimension 1 values (*r* =-.50, *p* =.006, FDR_adjusted_ *q* =.038) (see *Figure 4*). No significant correlations were detected between subdimension 1 values and EBA (*r* =-29, *p* =.130, FDR_adjusted_ *q* =.260), FBA (*r* =-.24, *p* =.212, FDR_adjusted_ *q* =.318), PMC (*r* =-.19, *p* =.342, FDR_adjusted_ *q* =.410), TPJ (*r* =-.03, *p* =.880, FDR_adjusted_ *q* =.599), or V1 (*r* =-.30, *p* =.126, FDR_adjusted_ *q* =.260) activity.

**Figure 4.**
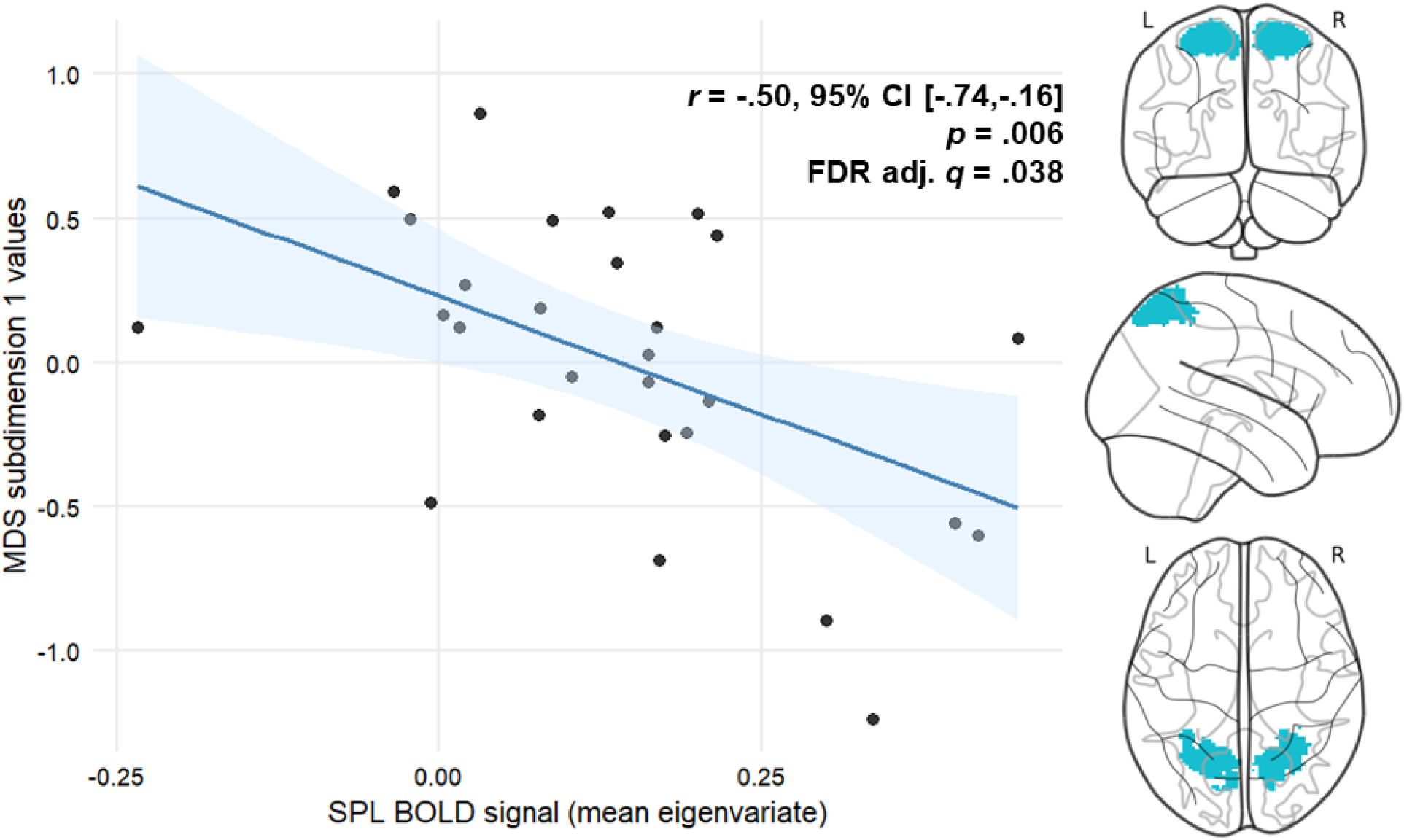
Scatterplot depicting a negative relationship between SPL mean BOLD signal (first eigenvariate) during BSE and MDS subdimension 1 values. On the right, three views of the SPL ROI from which the BOLD signal (averaged right and left) was extracted. Note. *MDS =* multidimensional scaling, *SPL* = superior parietal lobule, *BOLD* = blood-oxygen-level dependent, *L* = left, *R* = right.

### Exploratory Whole-brain Analyses

The whole-brain analysis for the avatar engagement contrast revealed significant activation across motor, visual, somatosensory, fronto-insular and cerebellar areas, spanning 9 clusters (see *Table 2* and *Figure 5*). Clusters were interpreted based on the Harvard-Oxford and Jülich Histological Atlas labels. The largest clustered in the right parietal operculum and extended dorsally into the superior and inferior parietal lobules. The second largest peak was observed in the right IFG and encompassed the ventral PMC, insula, and frontal operculum. A smaller cluster was identified in the same cortical territory in the left hemisphere. Bilateral occipitotemporal clusters were detected: one spanning right EBA, FBA, and another confined to left EBA. Activation was also observed in the medial superior frontal gyrus, including peaks in the supplementary motor area (SMA) and dorsal PMC. Additional clusters were detected in left IPL and left cerebellum, with peaks in lobule VI and lobule VIIb. No significant activity was associated with *BSE percent error*. Results from other event-related and parametric regressors are reported in the Supplement (see Supplementary Materials S4).

**Figure 5.**
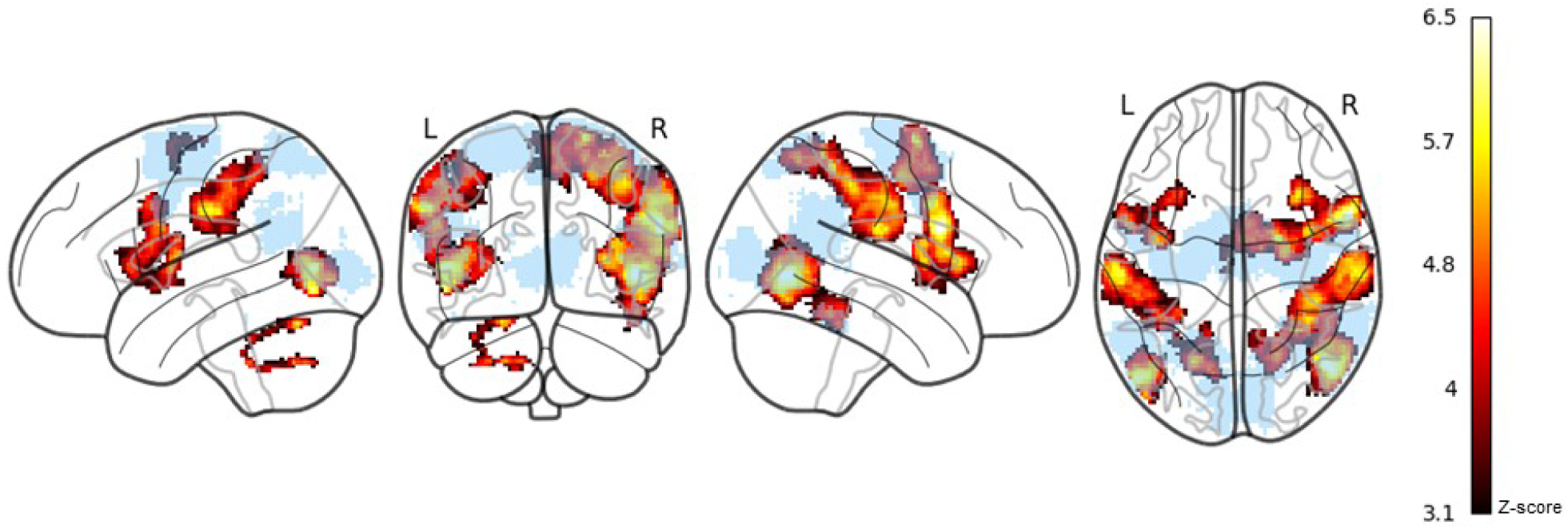
Statistical maps showing significant activation clusters from exploratory whole brain analysis. Regions showing increased BOLD response during avatar engagement, thresholded at *z* > 3.1. A dark red-to-yellow colour scale indicates increasing *z*-score magnitude. Maps are displayed on a glass brain. The ROI-mask (light blue) is overlaid to show regions in whole-brain analysis that were outside the a-priori mask.

**Table 2.** Clusters peaks associated with avatar engagement from whole-brain exploratory analyses.

| Cluster Peak | Size (voxels) | Z max | MNI Coordinates |  |  |
| --- | --- | --- | --- | --- | --- |
|  |  |  | *X* | *Y* | *Z* |
| R IPL/SPL | 2989 | 5.64 | 60 | -20 | 22 |
| R IFG | 1958 | 6.75 | 58 | 10 | 24 |
| R EBA | 1782 | 6.63 | 46 | -68 | 2 |
| R PMC | 1625 | 5.75 | 20 | -4 | 70 |
| L IPL | 1513 | 5.85 | -60 | -20 | 34 |
| L Precentral Gyrus | 1047 | 5.45 | -56 | 4 | 26 |
| L EBA | 697 | 6.16 | -50 | -74 | -4 |
| L Cerebellum – lobule VI | 225 | 5.49 | -22 | -66 | -24 |
| L Cerebellum – lobule VIIb | 126 | 5.06 | -20 | -64 | -44 |

*L* left hemisphere, *R* right hemisphere, *MNI* Montreal Neurological Institute, *IFG* inferior frontal gyrus, *IPL* inferior parietal lobule, *IPS* intraparietal sulcus, *MFG* middle frontal gyrus, *PMC* premotor cortex, *SMA* Supplementary Motor Area, *FG* fusiform gyrus, *EBA* Extrastriate Body Area. $Z > 3.1$ , cluster-corrected $p < .05$ FWE

## Discussion

This study aimed to characterize the neural activation patterns associated with BSE, and with BSE accuracy, in healthy individuals, using an interactive 3D avatar fMRI task. Consistent with our hypotheses, avatar engagement was associated with bilateral EBA, bilateral PMC, right FBA, and right SPL activation, but, contrary to our hypotheses, not TPJ or V1 activation. While no ROI, or region from the whole-brain analysis, showed significant modulation by trial-wise BSE accuracy, BOLD signal in the SPL was significantly associated with inter-individual differences in BSE error patterns, as indexed by MDS. Exploratory whole-brain analyses revealed broader task-related engagement across visuospatial, sensorimotor, and salience regions, and recapitulated the ROI results, demonstrating those regions’ robustness to whole-brain FWE correction. This study provides evidence for a functional map of brain regions engaged during whole, and body-part specific, size estimation, providing a reference point for future clinical research.

### Task-Evoked ROI Engagement

We hypothesized that six ROIs, previously implicated in visual and multisensory body representation, EBA, FBA, SPL, PMC, TPJ, and V1, would be activated during BSE-related engagement with the avatar. Four of these—bilateral EBA, right FBA, right SPL, and bilateral PMC—were significantly engaged. This activation pattern aligns with the task’s dual demands: participants needed to make localized adjustments to specific body parts while simultaneously maintaining a coherent global body configuration to accurately contextualize proportions of sizes and shapes. The EBA and FBA support part-based and whole-body form perception, respectively (Downing et al., 2001; Peelen & Downing, 2007; Taylor et al., 2007). The PMC, in the context of this task, may facilitate internal motor simulation during avatar manipulation (Hanakawa et al., 2008), while the SPL may support visuospatial mapping of body features (Medendorp & Heed, 2019).

By contrast, the TPJ and V1 were not significantly activated during the task. The TPJ, frequently implicated in perspective-taking, spatial self-location, and multisensory integration (Blanke et al., 2005; Tsakiris et al., 2007), may not have been substantially engaged due to the third-person (allocentric) vantage point from which participants viewed the avatar. Rather than adopting the avatar’s perspective (egocentric)—critical for evoking TPJ activity (Blanke et al., 2005; Martin et al., 2019, 2020; Wang et al., 2016), participants adjusted its form from an external viewpoint. The primary visual cortex processes the retinotopic and perceived size of objects, spatial frequency, and other low-level features such as contrast and orientation (Murray et al., 2006; Ng et al., 2006; Sperandio et al., 2012). While V1 likely contributes to initial size encoding, its activity may not have been prominent in the Somatomap 3D task, which emphasizes higher-order visual comparisons and spatial reasoning. These cognitive demands may be more effectively supported by downstream regions involved in body perception (EBA, FBA), visuospatial integration (SPL), and motor processes (PMC).

### Inter-Individual and Trial-Wise Associations Between BSE Accuracy and Neural Activity

A key finding was that SPL activity during BSE was significantly associated with inter-individual differences in BSE accuracy, as indexed by MDS. This result stands in contrast to the nonsignificant findings from parametric modulation analyses, which assess trial-by-trial fluctuations in BOLD signal relative to behavioral metrics. This suggests a dissociation between the neural correlates of stable, trait-like differences in body representation and the transient processing of accuracy at the level of individual body parts.

The SPL–MDS association reflects a between-subjects relationship: the degree to which an individual consistently engages the SPL during BSE relates to the structure of their estimation error profile across body parts. This could be due to the SPL’s involvement in global integrative functions— such as constructing and updating a spatial model of the body—rather than encoding transient estimation errors. Consistent with this role, the SPL integrates proprioceptive, tactile, and visual inputs (Schwoebel & Coslett, 2005) to maintain a dynamic, body-centered reference frame (Longo & Haggard, 2010; Zacks, 2008) and is implicated in spatial cognition and multisensory body schema (Medendorp & Heed, 2019). Within the context of Somatomap, SPL engagement may reflect efforts to align internally stored body representations with visual feedback from the manipulated avatar (Blanke et al., 2015).

The absence of significant modulation by trial-wise BSE accuracy may be due to the heterogeneity of neural processing across different body parts. Prior studies suggest that distinct cortical territories represent specific body parts in a topographically organized manner (e.g., the somatosensory homunculi), and even body-selective visual regions may exhibit differential sensitivity depending on body part identity (Bracci et al., 2015; Orlov et al., 2010). Thus, treating estimation accuracy across all body parts as repeated instances of the same phenomenon may have obscured region-or body-part-specific relationships. An alternative, future design could involve presenting multiple trials for each body part to allow body-part-specific modeling of accuracy-related signal, enabling more precise tests of regionally distinct neural contributions.

Notably, none of the other regions found to demonstrate significant task engagement— EBA, FBA, or PMC—showed significant correlations with individual differences in perceptual accuracy. This suggests that while these regions contribute to shared perceptual and motor imagery processes required during avatar engagement, they may not process stable, individualized representations of body size in the same manner as SPL.

### Exploratory Whole-Brain Analyses

Whole-brain analyses suggest engagement of distributed networks and regions. These included components of a visuospatial network (e.g., SPL) (Whitlock, 2017), components of a sensorimotor network (e.g., PMC, cerebellum) (Hanakawa et al., 2008; Stoodley et al., 2012), body-selective visual regions (e.g., EBA, FBA) (Taylor et al., 2007; Peelen & Downing, 2007), and components of the salience network (e.g., insula, frontal operculum) (Schimmelpfennig et al., 2023). Several of these regions—including the SPL, PMC, EBA, and FBA—overlapped with predefined ROIs, lending convergent support for their involvement in avatar engagement during the task.

Widespread activation was observed across parietal and frontal regions associated with both visuospatial and sensorimotor networks. Visuospatial regions included IPL, SPL, and postcentral gyrus—components of the multisensory body schema network, theorized to be involved in integrating proprioceptive, tactile, and visual input (Medendorp & Heed, 2019).

Sensorimotor-related activation included the dorsal PMC and SMA, regions associated with internally guided motor simulation and visuomotor transformation (Haggard, 2008; Hanakawa et al., 2008; Nachev et al., 2008). Posterior cerebellar lobules VI and VIIb were also engaged, consistent with their roles in sensorimotor coordination and cross-network integration involving attention and visual processing (Ji et al., 2019; Xue et al., 2021).

Body-selective and higher-order visual areas were engaged, including bilateral EBA, and right-lateralized FBA, largely overlapping with ROI analyses.

Lastly, activation was observed in regions commonly associated with salience-related processing, including the insula and frontal operculum. The insula, particularly its anterior portion, supports interoceptive awareness and affective salience (Craig, 2009; Simmons et al., 2013), while the adjacent frontal operculum has been identified in sustained vigilance and monitoring (Sadaghiani et al., 2010). These areas may support self-monitoring processes engaged during avatar adjustment, such as evaluating the match between visual input and internal body representation.

Taken together, from these exploratory results, we might speculate that avatar engagement recruits a distributed network that integrates perceptual, spatial, motor, and self-monitoring processes. While interpretations should remain cautious due to the exploratory nature of these analyses, the overlap with key ROI findings—particularly in the SPL, PMC, and EBA— provides convergent support for the core regions implicated in body estimation.

### Somatomap Task Performance

Participants showed systematic variation in both the magnitude and direction of estimation error across body parts. Percent error tended to be positive, indicating a general tendency to overestimate body part sizes, with the most pronounced overestimation observed for upper arm girth. However, consistent underestimation was observed for regions near the midsection, including torso length, abdomen protrusion, waist size, and hip size. This diverges from prior findings in clinical populations. For example, individuals with anorexia nervosa have demonstrated marked overestimation of midsection size using an earlier version of the Somatomap tool (Ralph-Nearman et al., 2019). In contrast, both Ralph-Nearman et al. and Karsan et al. (2024) found that healthy controls tended to underestimate waist and hip size while overestimating torso length. Thus, underestimation of waist and hip size appears to be a consistent finding across non-clinical samples.

Several methodological factors may account for these differences in body part estimation accuracy across studies that used Somatomap. First, the present study examined percent error, rather than centimeter error, which may better reflect perceptual accuracy given that just-noticeable differences for size scale proportionally with the magnitude of the stimulus (Fechner, 1966). Second, our study used an updated avatar that is smoother, has higher resolution, is more realistic, and has 26 adjustable body parts compared with 23 in earlier versions—including separately defined hip size and hip width. Additionally, the previous Somatomap 3D but not the current version matched skin and hair colour. Presenting a more minimal and abstract representation of the body, as in the previous Somatomap avatar, may alter reference cues used during estimation.

On average, participants spent approximately 3.3 seconds adjusting each body part, with interstimulus intervals averaging 4.1 seconds (see Supplementary Figure S5 for full distributions). These durations suggest that participants had sufficient time for deliberate perceptual judgments. These patterns suggest that healthy individuals make body-part-specific adjustments that likely reflect both perceptual tendencies—for example, greater salience or uncertainty around the torso—and structural features of the task, such as its top-down progression from head to feet.

### Limitations, Future Directions, and Clinical Implications

Several limitations in this study should be noted. First, the Somatomap 3D fMRI task was adapted from the previously-developed non-fMRI behavioural task, whose utility in quantifying BSE accuracy had been demonstrated in non-clinical samples (fashion models and non-fashion model controls (Ralph-Nearman et al., 2019, as proof-of-concept) and had been shown to have sensitivity to detect patterns of aberrant BSE accuracy in individuals with anorexia nervosa (Ralph-Nearman et al., 2021) and body dysmorphic disorder (Karsan et al., 2024) compared with healthy controls. In retaining most of the structure of the original task, it remains a continuous task (rather than having experiment-set discrete trials) and is self-paced. Yet, this partially unconstrained structure reduces experimental control and limits the ability to isolate discrete cognitive processes, e.g., perception vs. decision-making, which are likely intermingled during each body part trial. However, we partially mitigated concerns about variable trial and rotation durations and by using trial-wise response time and rotation duration as regressors. This allowed us to account for time-on-task effects and better isolate signal associated with estimation accuracy. Second, the relatively modest sample size (N = 28) may have limited statistical power. While our analytic approach addressed sources of variability, the sample size may have constrained sensitivity to detect smaller magnitude effects. Third, we applied a hypothesis-driven ROI approach, using several regions for which there was a priori knowledge of candidate body processing functions, but this may have overlooked other involved areas since the functional neural correlates of BSE, specifically, have not previously been well-established.

In terms of potential clinical applications, results from this study suggest that regions including the SPL, FBA, EBA, and PMC could be of interest in investigating neurobiological substrates of body image disturbances in those with eating disorders and body dysmorphic disorder. Among these, the SPL stands out for potential clinical relevance as its activity was associated with individual differences in BSE error profiles. This aligns with prior work implicating the posterior parietal cortex in body image distortion in anorexia nervosa (Gaudio & Quattrocchi, 2012; Mohr et al., 2010). Future studies could build on these findings by having individuals with anorexia nervosa complete this version of the Somatomap 3D task during fMRI. Alternatively, variations of the task could be developed to help discern specific cognitive processes. For instance, one variant could present brief, experimenter-controlled trials with fixed durations to better isolate perceptual and decision-related components. Outcomes such as atypical degree, spatial location, or timing of activation patterns, differences in sensitivity to perceptual uncertainty, or differences related to spatial transformation demands could help clarify how body image disturbances emerge and persist.

Finally, future studies that characterize the neural correlates of BSE in large populations could help establish normative models, which aims to quantify individual deviations from typical brain function and that can be stratified by, e.g. demographic or body morphometric factors such as height and weight (Marquand et al., 2016). Just as pediatric growth charts provide benchmarks for physical development, normative neuroimaging models can situate individuals along distributions of task-evoked activation, connectivity, or representational structure. Applied to BSE, this approach could help identify neural signatures of distorted self-perception, support individualized clinical assessment, and inform personalized interventions designed to restore more normative internal body representations.

## Conclusions

This study used a novel 3D avatar task to probe functional activation patterns associated with BSE, identifying a distributed neural system including visual, parietal, premotor, and self-monitoring regions. The SPL emerged as a key region associated with inter-individual differences in estimation accuracy, which potentially could contribute to perceptual body image distortions, suggesting significance for future clinical investigations. These findings advance efforts toward understanding the brain basis of BSE and offer specific targets for investigating altered body representation in clinical populations characterized by body image disturbances.

## CRediT Authorship Contribution Statement

**Hayden J. Peel:** Formal analysis, Visualization, Writing – original draft, Writing – review & editing. **Joel P. Diaz-Fong:** Data curation, Formal analysis, Investigation, Visualization, Writing – original draft, Writing – review & editing. **Sameena Karsan:** Data curation, Writing – review & editing. **Rajay Kumar:** Software, Writing – review & editing. **Gerhard Hellemann:** Methodology, Writing – review & editing. **Jamie D. Feusner:** Conceptualization, Funding acquisition, Investigation, Methodology, Supervision, Writing – review & editing.

## Acknowledgements

We would like to thank Bea Calahong and Katalin Groe for their help collecting data for this study. This research was enabled in part by support provided by Compute Ontario (<u>computeontario.ca</u>) and the Digital Research Alliance of Canada (<u>alliancecan.ca</u>). The Software used in this research was created by Jamie D. Feusner, Armen C, Arevian, and Nanthia Suthana (UCLA); and Sahib S. Khalsa and Christina Ralph-Nearman (Laureate Institute for Brain Research). © 2023 UCLA. Originally published in JMIR Mental Health (http://mental.jmir.org), 29.10.2019. Ralph-Nearman, C., Arevian, A. C., Puhl, M., Kumar, R., Villaroman, D., Suthana, N., Feusner, J. D., & Khalsa, S. S. (2019). A Novel Mobile Tool (Somatomap) to Assess Body Image Perception Pilot Tested With Fashion Models and Nonmodels: Cross-Sectional Study.

JMIR mental health, 6(10), e14115. https://doi.org/10.2196/14115

