## Supplementary Materials for "Functional Brain Mapping of Body Size Estimation Using a 3D Avatar"

**Author Note**

Hayden J. Peel, https://orcid.org/0000-0001-6078-2970
Joel P. Diaz-Fong, https://orcid.org/0000-0002-2108-6830
Sameena Karsan, https://orcid.org/0000-0003-0021-4810
Rajay Kumar, https://orcid.org/0000-0002-7306-4131
Gerhard Hellemann, https://orcid.org/0000-0003-2449-7704
Jamie D. Feusner, https://orcid.org/0000-0002-0391-345X

**Correspondence**

Jamie D. Feusner, 250 College St. #645, Toronto, Ontario M5T 1R8, Canada.


**Contents:**

**Supplementary Methods and Results**

1. Somatomap task details
2. Details on fMRIprep preprocessing
3. Behavioural performance during Somatomap
4. Other event and parametric modulation results

**Supplementary Figures**

**Figure S1** Mean response time per body part during body size estimation trials

**Figure S2** Distribution of body size estimation trial durations across all participants

**Figure S3** Distribution of avatar rotation durations during the task

**Figure S4** Distribution of inter-stimulus interval durations during the task

**Figure S5** Mean total time spent adjusting each body part during the task

**Figure S6** Stress plots for multidimensional scaling solutions across dimensionalities

**Figure S7** Heatmap showing Pearson correlation coefficients between estimation errors for individual body parts and scores on each of three subdimensions derived from multidimensional scaling

**Supplementary References**

1. Somatomap task details

*Somatomap avatar:* The 3D avatars used in this study were created from a male and a female human volunteer using a custom Raspberry Pi-based camera array consisting of 38 synchronized cameras. The resulting image sets were processed with RealityCapture software (v1.3; Epic Games, Inc., Cary, NC, United States) to generate high-resolution 3D body meshes. These meshes were then imported individually into Autodesk 3ds Max (Autodesk, Inc., San Rafael, CA, United States), where they were cleaned and smoothed into fully modifiable models. Custom controls were added to allow real-time adjustments of body part dimensions during the task.

*Randomization of body part parameters:* To mitigate potential anchoring effects, the initial positions of the size adjustment sliders were randomized for each body part. Specifically, the midpoint of each slider was jittered within ±10% of the true midpoint value prior to each trial. This ensured variability in the starting positions while preserving the full range of possible adjustments.

*fMRI task instructions:* "You will see an avatar with different body parts of random sizes. Your job is to adjust each body part to match your own. Use the right arrow to move through the parts one by one, adjusting sizes with the slider. To rotate the avatar horizontally, press and hold the left button and move the trackball; to move it vertically, use the right button. If you have difficulty navigating the avatar, hit 'Reset'. This will reset the position of the avatar, but not the sizes of the body parts. After adjusting all parts, you can go back to make additional adjustments. You have 10 minutes, but you can close your eyes if you finish early. Wait for our cue to begin."

*Mouse events:* The WebLink software (v2.2.161; SR Research Ltd., Ottowa, ON, Canada) was utilized to capture screen recordings, eye movements, and mouse and keyboard events, including the fMRI triggers used to synchronize events with the scan. Using mouse click timestamps, body size estimation was defined in the time-series as the periods when the participant was adjusting the avatar body size. Rotations were defined in the time-series as periods when the participant was rotating the position of the avatar. The baseline contrast included intervening periods between these rotations/adjustments.

Consecutive BSE events occurring less than one second apart were merged and treated as a single event. Additionally, all BSE and Rotation events lasting less than 200 milliseconds were discarded, as these were likely accidental or task-irrelevant clicks. BSE duration was defined as the time from the initial mouse click on the body part slider to the release of the mouse button, capturing the total interaction time per adjustment.

2. Details on fMRIprep preprocessing

Results included in this manuscript come from preprocessing performed using *fMRIPrep* 22.0.2 (Esteban, Markiewicz, et al. (2018); Esteban, Blair, et al. (2018); RRID:SCR_016216), which is based on *Nipype* 1.8.5 (K. Gorgolewski et al. (2011); K. J. Gorgolewski et al. (2018); RRID:SCR_002502).

*Preprocessing of $B_0$ inhomogeneity mappings*

A $B_0$-nonuniformity map (or *fieldmap*) was estimated based on two (or more) echo-planar imaging (EPI) references with topup (Andersson et al., 2003); FSL 6.0.5.1:57b01774).

*Anatomical data preprocessing*

A total of 1 T1-weighted (T1w) images were found within the input BIDS dataset. The T1-weighted (T1w) image was corrected for intensity non-uniformity (INU) with N4BiasFieldCorrection (Tustison et al. 2010), distributed with ANTs 2.3.3 (Avants et al. 2008, RRID:SCR_004757), and used as T1w-reference throughout the workflow. The T1w-reference was then skull-stripped with a *Nipype* implementation of the antsBrainExtraction.sh workflow (from ANTs), using OASIS30ANTs as target template. Brain tissue segmentation of cerebrospinal fluid (CSF), white-matter (WM) and gray-matter (GM) was performed on the brain-extracted T1w using fast (FSL 6.0.5.1:57b01774, RRID:SCR_002823, Zhang, Brady, and Smith 2001). Brain surfaces were reconstructed using recon-all (FreeSurfer 7.2.0, RRID:SCR_001847, Dale, Fischl, and Sereno 1999), and the brain mask estimated previously was refined with a custom variation of the method to reconcile ANTs-derived and FreeSurfer-derived segmentations of the cortical gray-matter of Mindboggle (RRID:SCR_002438, Klein et al. 2017). Volume-based spatial normalization to two standard spaces (MNI152NLin6Asym, MNI152NLin2009cAsym) was performed through nonlinear registration with antsRegistration (ANTs 2.3.3), using brain-extracted versions of both T1w reference and the T1w template. The following templates were selected for spatial normalization: *FSL's MNI ICBM 152 non-linear 6th Generation Asymmetric Average Brain Stereotaxic Registration Model* [Evans et al. (2012), RRID:SCR_002823; TemplateFlow ID: MNI152NLin6Asym], *ICBM 152 Nonlinear Asymmetrical template version 2009c* [Fonov et al. (2009), RRID:SCR_008796; TemplateFlow ID: MNI152NLin2009cAsym].

*Functional data preprocessing*

For each of the BOLD runs found per subject (across all tasks and sessions), the following preprocessing was performed. First, a reference volume and its skull-stripped version were generated by aligning and averaging 1 single-band references (SBRefs). Head-motion parameters with respect to the BOLD reference (transformation matrices, and six corresponding rotation and translation parameters) are estimated before any spatiotemporal filtering using mcflirt (FSL

6.0.5.1:57b01774, Jenkinson et al. 2002). The estimated *fieldmap* was then aligned with rigid-registration to the target EPI (echo-planar imaging) reference run. The field coefficients were mapped on to the reference EPI using the transform. BOLD runs were slice-time corrected to 0.452s (0.5 of slice acquisition range 0s-0.905s) using 3dTshift from AFNI (Cox and Hyde 1997, RRID:SCR_005927). The BOLD reference was then co-registered to the T1w reference using bbregister (FreeSurfer) which implements boundary-based registration (Greve and Fischl 2009). Co-registration was configured with six degrees of freedom. First, a reference volume and its skull-stripped version were generated using a custom methodology of *fMRIPrep*. Several confounding time-series were calculated based on the *preprocessed BOLD*: framewise displacement (FD), DVARS and three region-wise global signals. FD was computed using two formulations following Power (absolute sum of relative motions, Power et al. (2014)) and Jenkinson (relative root mean square displacement between affines, Jenkinson et al. (2002)). FD and DVARS are calculated for each functional run, both using their implementations in *Nipype* (following the definitions by Power et al. 2014). The three global signals are extracted within the CSF, the WM, and the whole-brain masks. Additionally, a set of physiological regressors were extracted to allow for component-based noise correction (*CompCor*, Behzadi et al. 2007). Principal components are estimated after high-pass filtering the *preprocessed BOLD* time-series (using a discrete cosine filter with 128s cut-off) for the two *CompCor* variants: temporal (tCompCor) and anatomical (aCompCor). tCompCor components are then calculated from the top 2% variable voxels within the brain mask. For aCompCor, three probabilistic masks (CSF, WM and combined CSF+WM) are generated in anatomical space. The implementation differs from that of Behzadi et al. in that instead of eroding the masks by 2 pixels on BOLD space, a mask of pixels that likely contain a volume fraction of GM is subtracted from the aCompCor masks. This mask is obtained by dilating a GM mask extracted from the FreeSurfer's *aseg* segmentation, and it ensures components are not extracted from voxels containing a minimal fraction of GM. Finally, these masks are resampled into BOLD space and binarized by thresholding at 0.99 (as in the original implementation). Components are also calculated separately within the WM and CSF masks. For each CompCor decomposition, the *k* components with the largest singular values are retained, such that the retained components' time series are sufficient to explain 50 percent of variance across the nuisance mask (CSF, WM, combined, or temporal). The remaining components are dropped from consideration. The head-motion estimates calculated in the correction step were also placed within the corresponding confounds file. The confound time series derived from head motion estimates and global signals were expanded with the inclusion of temporal derivatives and quadratic terms for each (Satterthwaite et al. 2013). Frames that exceeded a threshold of 0.5 mm FD or 1.5 standardized DVARS were annotated as motion outliers. Additional nuisance timeseries are calculated by means of principal components analysis of the signal found within a thin band (*crown*) of voxels around the edge of the brain, as proposed by (. The BOLD time-series were resampled into standard space, generating a *preprocessed BOLD run in MNI152NLin6Asym space*. First, a reference volume and its skull-stripped version were generated using a custom methodology of *fMRIPrep*. Automatic removal of motion artifacts using independent component analysis (ICA-AROMA, (Pruim et al., 2015) was performed on the *preprocessed BOLD on MNI space* time-series after removal of non-steady state volumes and spatial smoothing with an isotropic, Gaussian kernel of 6mm FWHM (full-width half-maximum). Corresponding "non-aggressively" denoised runs were produced after such smoothing. Additionally, the "aggressive" noise-regressors were collected and placed in the corresponding confounds file. All resamplings can be performed with *a single interpolation step* by composing all the pertinent transformations (i.e. head-motion transform matrices, susceptibility distortion correction when available, and co-registrations to anatomical and output spaces). Gridded (volumetric) resamplings were performed using antsApplyTransforms (ANTs), configured with Lanczos interpolation to minimize the smoothing effects of other kernels (Lanczos 1964). Non-gridded (surface) resamplings were performed using mri_vol2surf (FreeSurfer).

### 3. Behavioural performance during Somatomap

Mean response times varied across body parts (see *Supplementary Figure* S1), with the longest durations observed for adjustments to the torso and midsection (e.g., abdomen protrusion, torso length, waist size), while areas such as the feet and ankles showed the shortest response times.

Participants typically spent more time on adjustments early in the task sequence (i.e., towards the top of the body) and became faster as the task progressed, with peaks in RT centered on midline regions like the torso and waist. Notably, the task allowed for free exploration: participants could revisit and re-adjust body parts multiple times. As such, these response times likely reflect a blend of initial decision difficulty, perceptual uncertainty, and iterative refinement, rather than isolated judgements.

Note, both BSE size adjustment durations and rotation durations were positively skewed, as is common in response time data. However, we did not apply any transformations to these durations, as doing so would distort the actual temporal structure of the events as they occurred during scanning. Instead, we addressed this skew statistically by modeling duration-related variability using separate parametric regressors for BSE and rotation events in the general linear model. This allowed us to account for the influence of trial-wise duration on neural activity while preserving the integrity of the original event timing.

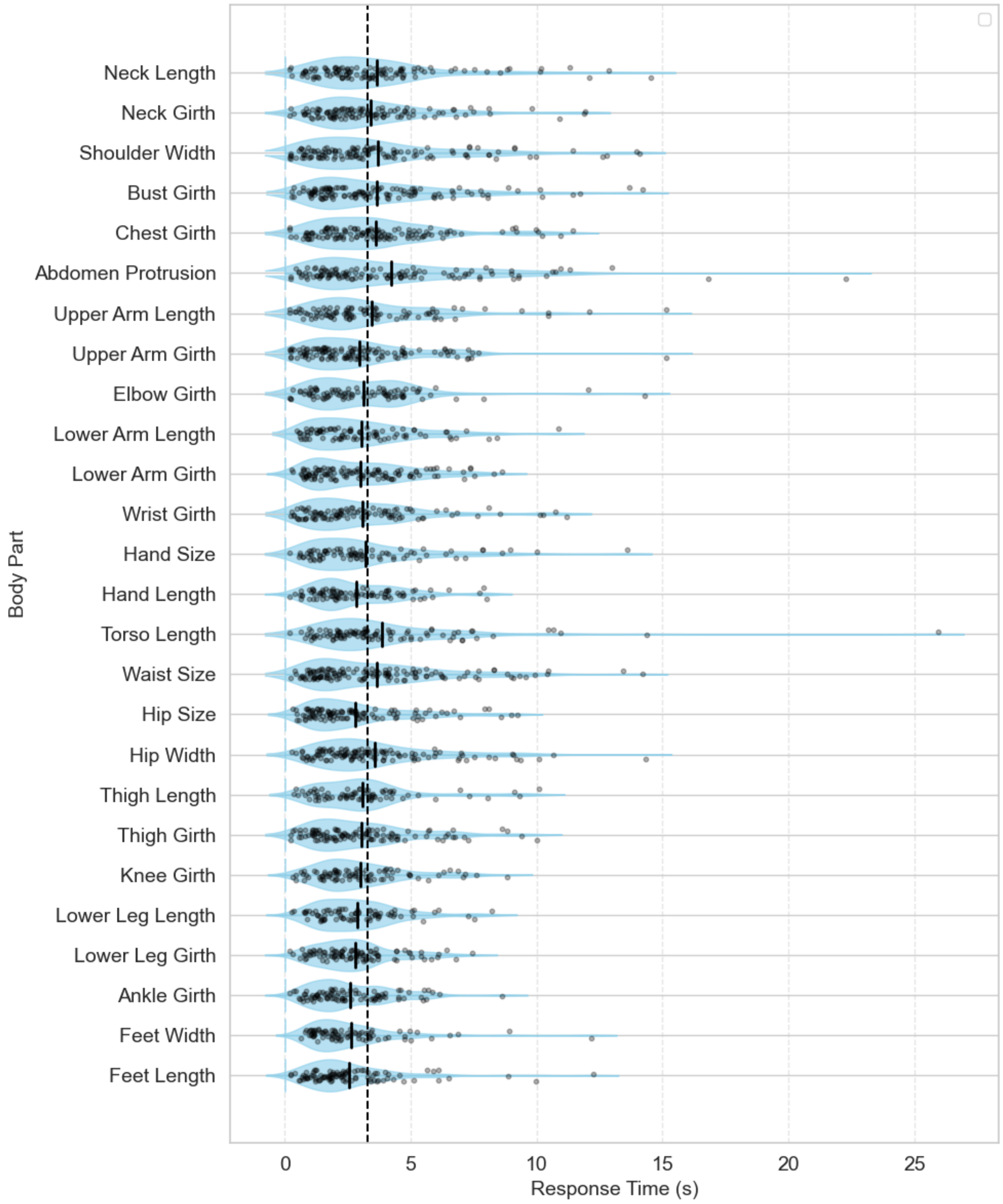


*Supplementary Figure S1*. Mean response time (RT) per body part during body size estimation (BSE) trials. Body parts are ordered anatomically from head (top) to feet (bottom). Bars indicate the average RT in seconds, with error bars reflecting the standard error of the mean. A vertical dashed line indicates the grand mean RT across all BSE trials (3.28 s).

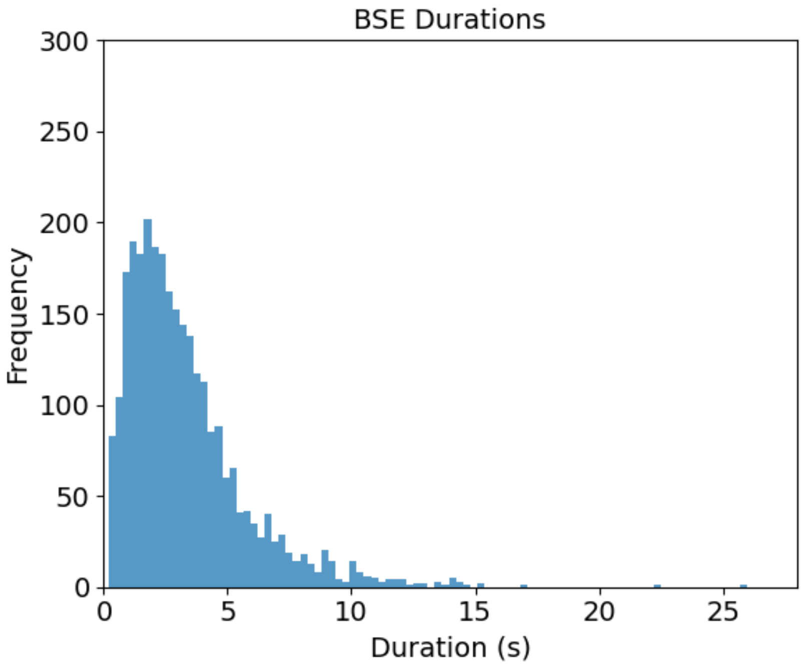


*Supplementary Figure S2.* Distribution of body size estimation (BSE) trial durations across all participants. Histogram displays the frequency of trials as a function of response duration (in seconds), reflecting the time participants spent adjusting each body part. Durations are pooled across body regions and participants.

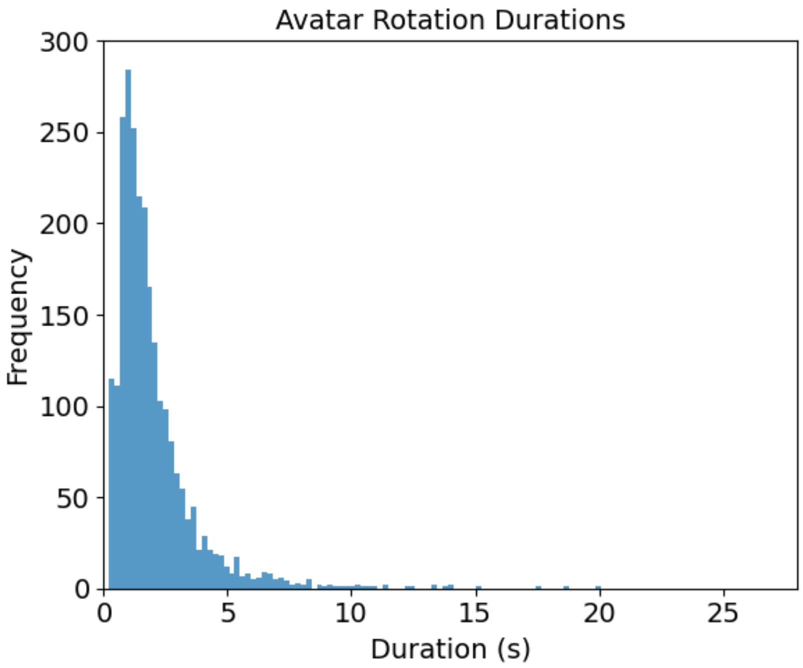


*Supplementary Figure S3.* Distribution of avatar rotation durations during the task. The histogram depicts the frequency of rotation events as a function of duration (in seconds), aggregated across all participants. Rotation durations reflect the time spent manipulating the avatar's orientation between body part adjustments.

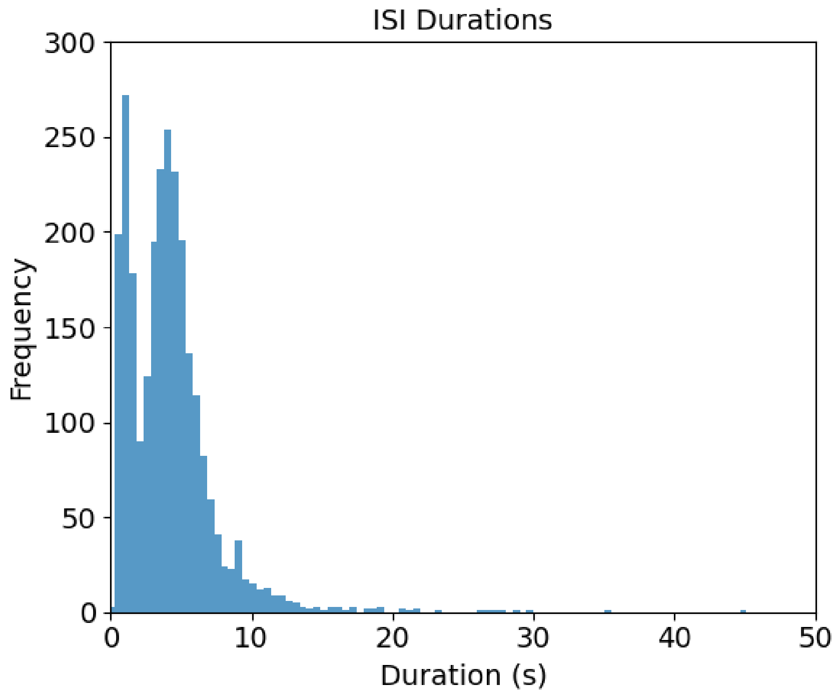


*Supplementary Figure S4.* Distribution of inter-stimulus interval (ISI) durations during the task. The histogram shows the frequency of ISI events as a function of duration (in seconds), aggregated across all participants. ISIs reflect periods between discrete task events, such as transitions between rotation and size adjustment phases. The observed bimodal distribution likely reflects two distinct classes of ISIs: shorter durations between BSE estimations or between BSE and rotations within the same body part, and longer durations associated with navigating between different body parts (e.g., navigating to the "next" button and returning to the main interface).

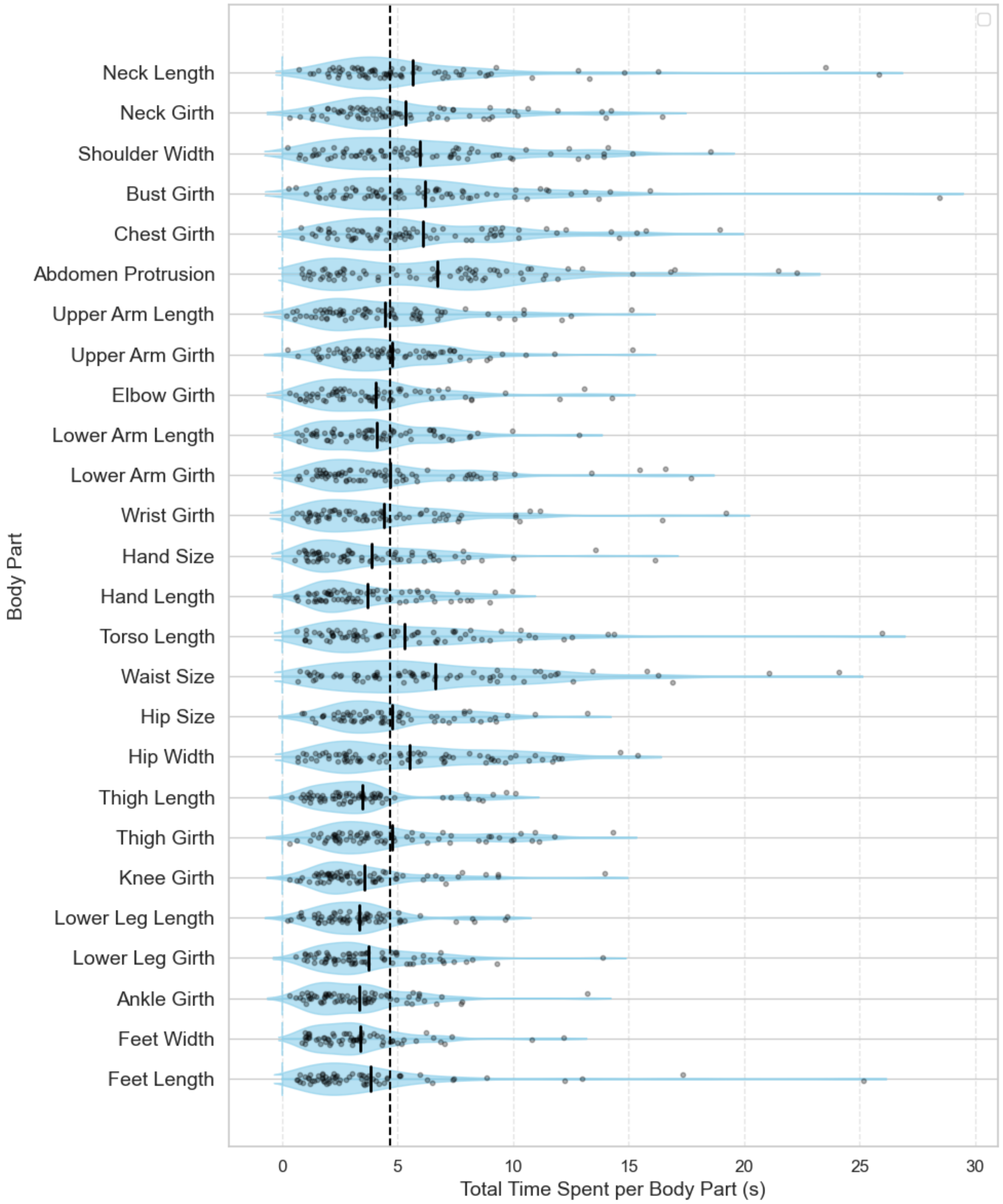


*Supplementary Figure S5*. Mean total time spent adjusting each body part during the task. Bars represent the average cumulative duration (in seconds) per body part across all participants. Error bars indicate ±1 standard error of the mean.

4. Other event and parametric modulation results

In addition to primary regressors of interest, other regressors were included when constructing the model. These included separate BSE size adjustment and rotation events (to construct the 'avatar engagement' contrast), as well as parametric regressors for rotation events, BSE RT, and rotation RT.

*BSE size adjustment events*

*Within-mask analyses:* BSE size adjustment events were associated with activity in 4 clusters: right EBA (cluster size = 339, Z = 5.72, peak in MNI space = 46 –66 –4), left EBA (cluster size = 196, Z = 5.69, peak in MNI space = -50 -72 –2), right FBA (cluster size = 40, Z = 4.52, peak in MNI space = 44 –48 -16), and right PMC (cluster size = 40, Z = 4.04, peak in MNI space = 6 0 66)

*Whole brain analyses:* For whole brain analyses, BSE size adjustment events were associated with 10 clusters of activity. This included the right precentral gyrus (cluster size = 1,650, Z = 6.44, peak in MNI space = 52 10 24), right inferior parietal lobule (IPL; cluster size = 1,519, Z = 5.58, peak = 60 –26 40), right extrastriate body area (EBA; cluster size = 1,029, Z = 5.72, peak = 44 –66 –4), left EBA (cluster size = 396, Z = 5.69, peak = –50 –72 –2), left IPL (cluster size = 215, Z = 5.22, peak = –58 –24 32), left insula (cluster size = 195, Z = 4.75, peak = –4 4 10), right frontal pole (cluster size = 168, Z = 4.31, peak = 10 42 10), right frontal orbital cortex (cluster size = 82, Z = 4.67, peak = 26 30 –10), left insula (cluster size = 74, Z = 4.36, peak = –30 –32 4), and left inferior frontal gyrus (IFG; cluster size = 65, Z = 4.47, peak = –56 10 14).

*Rotation events*

*ROI analyses:* Rotation events were associated with activity in 7 clusters: left PMC (cluster size = 2299, Z = 5.91, peak in MNI space = 26 –12 54), left SPL (cluster size = 596, Z = 5.39, peak in MNI space = -14 –60 62), right SPL (cluster size = 544, Z = 6.41, peak in MNI space = 32 –50 62), right EBA (cluster size = 515, Z = 6.98, peak in MNI space = 46 –68 2), left EBA (cluster size = 274, Z = 6.16, peak in MNI space = -46 –74 4), right FBA (cluster size = 96, Z = 4.65, peak in MNI space = 44 –40 –18), left PMC (cluster size = 59, Z = 5.26, peak in MNI space = -60 6 32).

*Whole brain analyses:* For whole brain analyses, rotation events were associated with 11 clusters of activity. This included bilateral SPL (cluster size = 15,661, Z = 6.41, peak in MNI space = 32 –50 62), right EBA (cluster size = 2,752, Z = 6.98, peak = 46 –68 2), frontal operculum / insula (cluster size = 2,752, Z = 6.98, peak = 46 –68 2), left cerebellum (cluster size = 984, Z = 5.34, peak = –16 –58 –46), left EBA (cluster size = 895, Z = 6.16, peak = –46 –74 4), right thalamus (cluster size = 239, Z = 4.68, peak = –12 –12 8), left frontal pole (cluster size =

225, Z = 4.45, peak = –40 42 36), right middle frontal gyrus (cluster size = 197, Z = 4.30, peak = 30 34 24), left cingulate (cluster size = 195, Z = 4.22, peak = 24 –40 –13), posterior intraparietal sulcus (cluster size = 184, Z = 4.13, peak = 26 –82 44), and left pallidum (cluster size = 141, Z = 4.57, peak = –16 4 0).

*BSE size adjustment RT parametric modulation*

*Within-mask analyses:* Longer response times for BSE size adjustment events were associated with activity in the right SPL (cluster size = 237, Z = 5.05, peak in MNI space = 50 –44 20), left V1 (cluster size = 218, Z = 5.36, peak in MNI space = -10 –88 0), and left SPL (cluster size = 122, Z = 4.13, peak in MNI space = -56 –54 22, and cluster size = 38, Z = 4.79, peak in MNI space = -60 –52 38).

*Whole brain analyses:* For whole brain analyses, longer BSE response times were associated with 9 clusters. This included left primary visual cortex (V1; cluster size = 1,474, Z = 5.64, peak in MNI space = –12 –90 –10), right superior parietal lobule (SPL; cluster size = 1,087, Z = 5.05, peak = 50 –44 20), left SPL (cluster size = 887, Z = 4.79, peak = –30 –52 38), left inferior frontal gyrus (IFG; cluster size = 314, Z = 4.67, peak = –46 22 12), right middle frontal gyrus (cluster size = 233, Z = 4.61, peak = 44 16 32), right V1 (cluster size = 194, Z = 4.17, peak = 22 –98 4), left precuneus (cluster size = 144, Z = 4.26, peak = –4 –56 52), right cerebellum (cluster size = 141, Z = 4.24, peak = 4 –80 –30), and left IFG (cluster size = 103, Z = 4.67, peak = –44 18 26).

*Rotation RT parametric modulation*

*Within-mask analyses:* Longer response times for rotation events were not associated with any activity in ROIs.

*Whole brain analyses:* For whole brain analyses, there were again no effects detected for rotation RT.

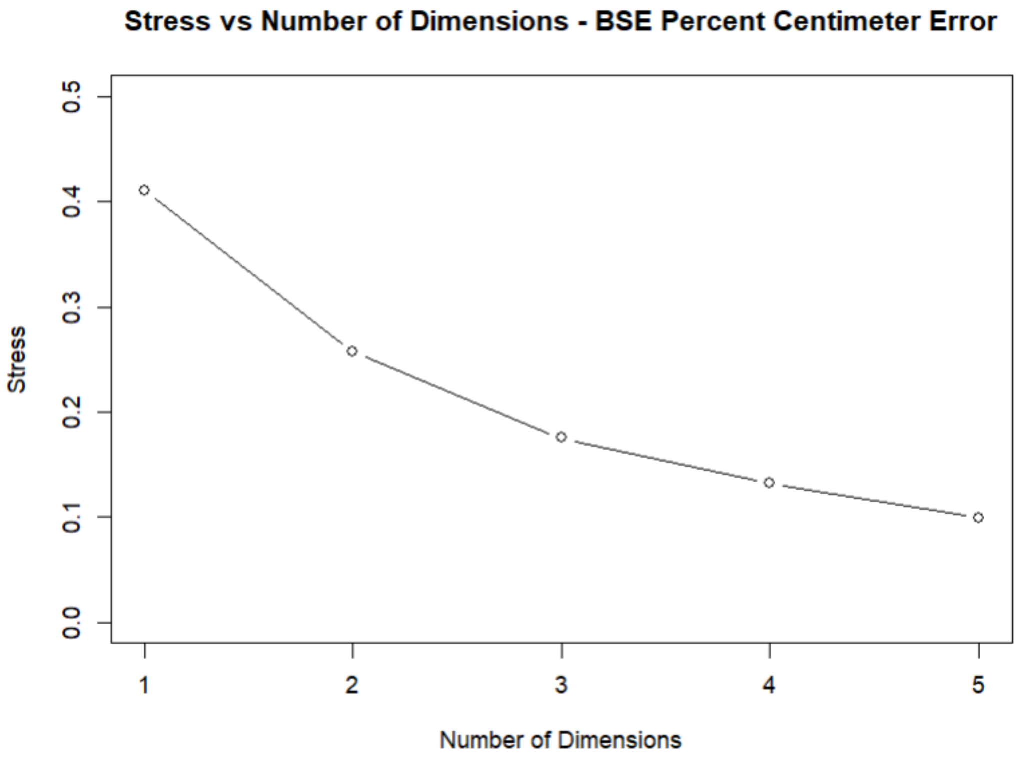


*Supplementary Figure S6.* Stress plots for multidimensional scaling (MDS) solutions across dimensionalities. Plots depict the relationship between the number of retained dimensions and Kruskal's stress values, used to assess goodness-of-fit.

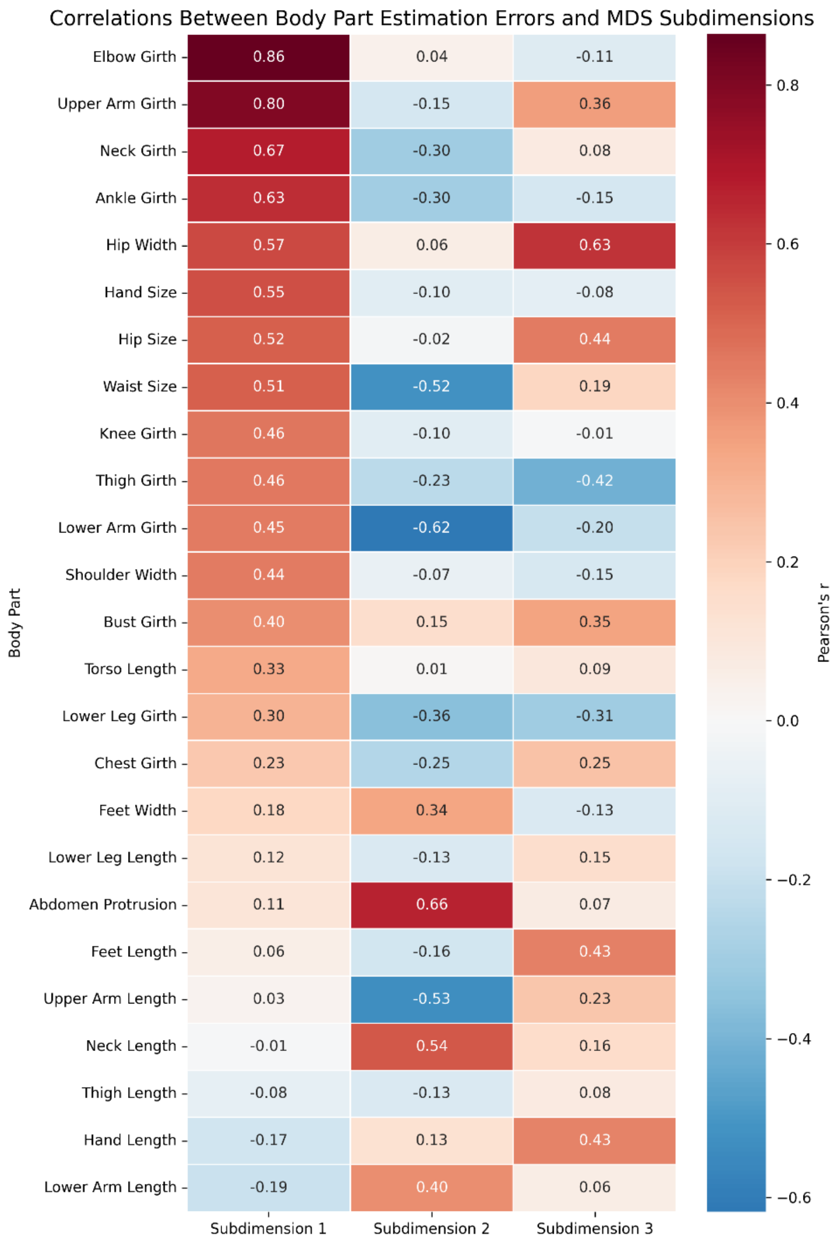


*Supplementary Figure S7.* Heatmap showing Pearson correlation coefficients (r) between estimation errors for individual body parts and scores on each of three subdimensions derived from multidimensional scaling (MDS). Warmer colors indicate positive correlations, and cooler colors indicate negative correlations.
